# Dormancy Breaking Treatments in Northern Wild Rice (*Zizania palustris* L.) Seed Suggest a Physiological Source of Dormancy

**DOI:** 10.1101/2021.09.10.459785

**Authors:** Lillian McGilp, Aaron Semington, Jennifer Kimball

## Abstract

Dormancy is a limiting factor for breeding in northern wild rice (NWR; *Zizania palustris* L). This study developed a dormancy curve and tested a combination of scarification and hormone treatments, across three timepoints, for their ability to break dormancy in NWR and produce viable seedlings and plants. A dormancy curve was established across 9 months post-harvest, which showed maximum germination (95%) by 17 weeks post-harvest and high germination (≥81 %) through the rest of the testing period. Next, dormancy breaking treatments were tested. At 1 week post-harvest, few seeds germinated (≤ 15 %) across all treatment combinations. However, sulfuric acid increased germination shortly after harvest (5.8 %), compared to water (0.5 %) and NaClO (0 %) but resulted in stunted seedlings, all but one of which died shortly thereafter. At 7 weeks, sulfuric acid treated seeds did not result in significantly higher germination than water and maximum germination was still below 15%. By 11 weeks post-harvest, the water treatments had the highest germination and resulted in the most viable plants, indicating that dormancy had begun to break naturally and exceeded the effect of the other scarification treatments. Hormonal treatments had no significant effect on germination or seed viability and no strong conclusions could be drawn about their effect on seedling or plant health. Due to the inability of early germinated seed to consistently produce viable plants and the increase in germination following sufficient cold storage, it is likely that NWR seed has intermediate or deep physiological dormancy.

## Introduction

Northern wild rice (NWR; *Zizania palustris* L.) is an outcrossing, annual aquatic grass in the Poaceae family. The species grows naturally in lakes and streams across the Great Lakes region of the United States and Canada, and provides food and habitat to many species in aquatic ecosystems (Steeves, 1952; Huseby, 1997; Biesboer, 2019). In addition to natural stands, NWR has been cultivated in irrigated paddies since the 1950s, with the largest production in Minnesota and California (Oelke and Porter, 2016). NWRs unique seed physiology is a major challenge for the development of new cultivars, particularly due to the limited longevity of seed in current storage conditions. This loss of viability during NWR storage is related to the intermediately recalcitrant, or desiccation sensitive, seed, which is commonly associated with short-term viability and limits *ex-situ* seed storage (Probert and Brierley, 1989; Kovach and Bradford, 1992a; Dickie and Pritchard, 2002; McGilp *et al*., 2020). NWR seed also has a dormancy period of about three to six months, during which cool (1-3 °C), submerged, and dark conditions are necessary for the stratification of seed (Simpson, 1966; Cardwell *et al*., 1978; Grombacher *et al*., 1997). The dormancy period of NWR seed may initially extend the viability of seed, compared to other recalcitrant species, however, as dormancy breaks over this period, the seed begins to lose viability quickly, preventing the long-term use of germplasm. Additionally, the dormancy period in NWR limits the number of cycles of selection possible per year, slowing the progress of breeding. Although both recalcitrance and dormancy can be limiting factors in the breeding of NWR, little is known about the mechanisms underlying these traits or the relationship between them. This study aims to increase the knowledge about the basic biology underlying dormancy in NWR.

Seed dormancy is the absence of germination, within a certain period of time, despite the presence of favorable environmental conditions, including but not limited to, temperature, light quality, and the availability of light, water, and oxygen (Hilhorst, 1995; Bewley *et al*., 2013). While different types of dormancy have been defined in various ways over time, they can be broken down into 5 broad categories including: physical (PY), or dormancy related to impermeability of the seed coat, physiological (PD), where there is a block to the expansion of the embryo, morphological (MD), in which the embryo is undifferentiated or underdeveloped, morphophysiological (MPD), a combination of MD and PD, or combinational (PY + PD), a combination of PY and PD (Baskin and Baskin, 2004, 2014a). It is thought that in NWR seed there are multiple mechanisms for the control of dormancy (Oelke, 1982). The tough, waxy seed coat of NWR appears to block the embryo from expanding, either due to impermeability restricting the imbibition of seed or via physical restriction by the seed coat, which is evidenced by the increase in germination upon mechanical removal of the pericarp overlaying the embryo (Woods and Gutek, 1974; Cardwell *et al*., 1978; Oelke, 1982). Additionally, it is likely that there is hormonal inhibition of embryo growth in NWR as evidenced by large quantities of abscisic acid (ABA) found in dormant NWR seed and reduced germination of hulled seeds when suspended in an aqueous solution of hulls (Cardwell *et al*., 1978; Albrecht *et al*., 1979; Oelke and Albrecht, 1980; Grombacher *et al*., 1997). However, the specific type of dormancy in NWR has not been clearly defined.

In an effort to learn more about dormancy in NWR, as well as increase the number of possible breeding cycles in a year, several studies have evaluated methods of overcoming dormancy. Some research has focused on breaking dormancy by breaching the seed coat, including the physical removal of portions of the pericarp with a scalpel, dissecting needle, or through the use of abrasive granules, as well as the application of chemicals such as ethanol, chloroform, sodium hypochlorite, and acetone (Simpson, 1966; Oelke and Albrecht, 1978, 1980). Overall, these studies found only modest increases in the germination of NWR seed, which did not always correspond to higher seedling vigor. Other studies have tested the application of hormones to overcome seed dormancy in NWR. In particular, researchers have evaluated the efficacy of applications of gibberellic acid (GA_3_), 6-benzyl adenine (BA), and kinetin for the germination of dormant seed (Simpson, 1966; Cardwell *et al*., 1978; Oelke and Albrecht, 1980). As compared to treatments that were only scarified, treatments with GA_3_, especially when combined with kinetin, had a greater effect on the germination of NWR seed. Similar studies in white rice (*Oryza sativa* L.) have also demonstrated the efficacy of the application of kinetin and GA_3_ in overcoming seed dormancy (Cohn and Butera, 1982; Miyoshi and Sato, 1997; Vieira *et al*., 2002). While some NWR seed treatments have shown modest increases in germination, there is not yet a sufficiently efficacious method on which breeding programs can rely.

Despite the breeding challenges imposed by it, little is known about the dormancy of NWR seed, particularly compared to other cultivated plant species. Although previous studies have evaluated the efficacy of various treatments for breaking dormancy in NWR, many of them have used dehulled seed or seed with punctured pericarps, processes which are not feasible for large scale breeding and have resulted in weakened seedlings (Oelke and Albrecht, 1978, 1980). Additionally, only a handful of these studies have been conducted early in the dormancy period, during which maximal inhibition would be present. Finally, previous studies have not evaluated the use of sulfuric acid as a means of weakening the pericarp of NWR, despite its use in other species (Li *et al*., 1999; Purohit *et al*., 2015; Lata *et al*., 2018; Xiong *et al*., 2018). Therefore, a re-evaluation of dormancy breaking techniques for NWR is needed. The objectives of this study were to 1) Evaluate the efficacy of the combination of mechanical and hormonal treatments for early dormancy break in NWR and 2) Assess the effect of early dormancy break on the viability of the resulting NWR plants. In this study, mechanical and hormonal treatments were applied individually or successively to NWR seed, at three timepoints, throughout the initial stages of dormancy. The efficacy of the treatments was evaluated through the measurement of germination, metabolic activity, seedling size, and plant health.

## Materials and Methods

### Plant Materials

In 2019, the NWR cultivated variety ‘Itasca-C12’ and the breeding line ‘K2EF-C16’, representing genotypes with average and early germination, respectively, were used to establish a dormancy curve. Itasca-C12 was broadcast planted at a density of 112 kg ha^−1^, in a grower’s dike irrigated, field in Gully, MN. K2EF-C16 was broadcast planted at a density of 15.7 kg ha^−1^ in a 12 × 15 m, in a dike-lined research paddy at the University of Minnesota North Central Research and Outreach Center (NCROC), in Grand Rapids, MN (47.2372° N, 93.5302° W, and 392 m elevation). The genotypes were both planted on May 16^th^ of 2019, open pollinated in their respective paddies, and then hand harvested on September 3^rd^ and 4^th^ of 2019, respectively.

For the dormancy breaking experiments, the NWR variety ‘Barron’ was chosen, as it is a productive variety with good germination, and excess seed was readily available for the experiment. Germination was an important consideration for this study, as the recalcitrance of NWR seed limits the viability of seed stocks in storage. Seeds were tractor-planted planted on May 12^th^ of 2020, at 15.7 kg ha^−1^, in a 153 × 27 m dike irrigated research plot, with 3m single rows, spaced 0.38m apart, and 1m alleys, for a final plant density of 1 plant per 0.09m^2^. Plants were open pollinated and seed was hand harvested on September 10th of 2020. Following harvest and processing, seed was stored submerged in dH_2_0 in heavy sealed plastic bags, in the dark, at 3°C until the beginning of each trial.

### Germination and Tetrazolium Testing

To assess seed viability, seeds were placed in petri dishes lined with filter paper and 20 ml of dH_2_0. Plates were then randomized within blocks, in a Conviron-E15 growth chamber (Winnipeg, Canada), for germination testing. The growth chamber was set to 15-hour days at 20°C and 9-hour nights at 14°C. To keep seeds moist through-out the germination testing period, dH_2_0 was added to petri dishes as necessary. After 14 days, the number of germinated seeds, as defined by coleoptile emergence equal to the length of the seed, were quantified. The germinated seeds from the dormancy treatment experiment were then placed in 4 oz jars with 20 ml of dH_2_0 for 1 week, in preparation for seedling evaluations and remained in the growth chamber under the same conditions as for germination testing. dH_2_0 was added to jars as necessary to keep seeds submerged.

Ungerminated seeds were evaluated for metabolic activity to estimate seed viability with a protocol adapted from the tetrazolium testing handbook (Peters, 2000). Briefly, seed was bisected longitudinally to expose the embryo, then suspended in a 0.2% solution of 2,3,5 triphenyl tetrazolium chloride (Mp Biomedicals Inc, Santa Ana, CA) for 2 hours. Seed was then triple-rinsed and the number of stained embryos was quantified.

### Evaluation of NWR Dormancy

To produce a dormancy curve for NWR, seed was evaluated over a 9-month period (~38 weeks), starting from harvest, on September 5^th^, 2019, through May 27th, 2020. Every two weeks during this period, 3 replicates of 20 seeds were pulled from submerged cold storage. Seeds were germination tested, under the same conditions as is described above. Any seed that did not germinate, was subsequently tetrazolium stained, as described above, to assess seed viability. From these data, a dormancy curve was produced and an estimate of the weeks post-harvest at which 50 % of viable seed germinated was calculated, as described in the data analysis section.

### Dormancy Breaking Experiment

To develop improved methodology for overcoming dormancy in NWR, we evaluated several treatments, over the first three months of the dormancy period. The experiment was repeated 3 times, with the first experiment starting on September 21^st^, 2020, 1 week post-harvest, the second on October 27^th^, 7 weeks post-harvest, and the third on November 24^th^, 11 weeks post-harvest. The trials were split into five steps: primary treatment, targeting the seed coat, secondary treatment, using hormones to stimulate the embryo, germination and tetrazolium testing, seedling evaluations, and plant health ratings. Each trial was set up as a randomized complete block design, with 5 blocks of 20 seeds per primary and secondary treatment combination.

To evaluate the possibility of seed coat imposed dormancy, or that related to the tissues surrounding the embryo, three primary treatments were tested, including a 2 hour soak in 300 ml of 2.63 % sodium hypochlorite (NaClO) (Oelke and Albrecht, 1980), a 45 second wash in 200 ml of 98 % sulfuric acid (SA), and a 45 second wash in dH_2_0, as a control. Following the treatment of seeds with NaClO, SA, or water, seeds were rinsed 3 times with 300 ml of dH_2_0. To evaluate the potential for embryo imposed dormancy, or that related to the embryo or endosperm, secondary hormone treatments were immediately applied following primary treatments. Secondary treatments included: 0.1 mM gibberellic acid (GA_3_) (Research Products International, Mt Prospect, IL), 1 mM kinetin (K) (Chem Impex International Inc, Bensenville, IL), a combination of 0.1 mM GA_3_ and 1 mM K, and a dH_2_0 control. Seeds were placed in sealed, 4 oz plastic jars, with 20 ml of secondary treatment solution and stored at room temperature (~24°C), in the dark, for 24 hours (Oelke and Albrecht, 1980). After 24 hours, each jar of seed was rinsed for 30 seconds under running water.

### Evaluation of Seedling and Plant Vigor

To evaluate the seedling and plant vigor of seeds that germinated, seedlings and plants were assessed by several methods. After 1 week of growth in the growth chamber, seedlings were measured from root tip to the top of shoot tissue. Seedlings were then randomized in a greenhouse, set to 12-hour days at 24°C and 12-hour nights at 18°C. Seedlings were grown in fully submerged, 25 cm deep, 3.2 cm radius cone-tainers, amended with 22.4 kg ha^−1^ of urea (Alpha Chemicals, Stoughton, MA) and 48.8 kg ha^−1^ of Sprint 330 iron chelate (BASF, Ludwigshafen, Germany) for 30 days. After 30 days, the root and shoot tissue of each plant were measured separately, using a measuring tape, and plants were given a health rating of 1-4 (1: a dead plant, 2: a plant with significantly stunted growth and thin leaves, 3: a plant with relatively normal growth but with minor stunting of stem or leaf tissue, 4: A healthy plant with no stunting) (Figure 1).

**Figure 1.**
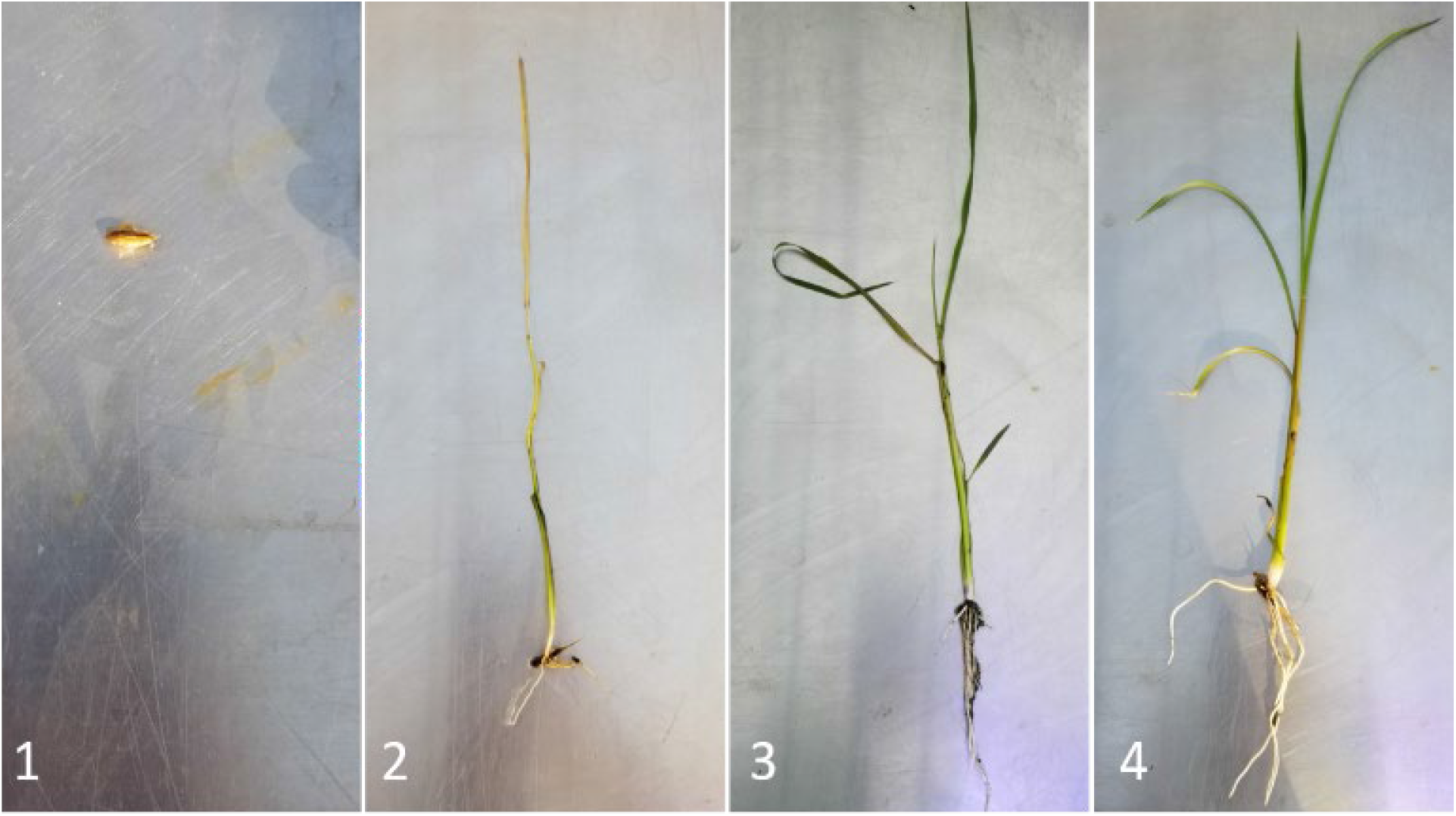
Plant health rating scale at 1 month of growth in the greenhouse of northern wild rice (NWR; *Zizania palustris* L.) following treatment with primary and secondary dormancy breaking treatments. 1: a dead plant, 2: a plant with significantly stunted growth and thin leaves, 3: a plant with relatively normal growth but with minor stunting of stem or leaf tissue, 4: A healthy plant with no stunting.

### Data Analysis

Analyses were done in Rstudio, version 1.2.5001 (Rstudio-team, 2019). Data was first run through standard residual diagnostic plots, including residuals versus fitted, normal Q-Q, scale-location and residuals versus leverage, to assess data quality. An analysis of variance (ANOVA) was conducted for germination and tetrazolium testing results. Significant results from the ANOVA were evaluated using the post hoc Tukey LSD test from the ‘agricolae’ package, using a Bonferroni adjustment for p-values (Mendiburu, 2019). Due to uneven sample sizes in the seedling and plant rating data, statistical analyses could not be done on these data. The ‘ggplot2’ package (Wikham, 2016) was used to produce all graphs in this publication.

The equation 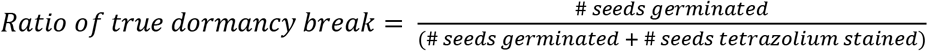 was used to determine the ratio of seeds that had germinated, per time point, relative to the number of tested seeds that were actually viable. A modified LD_50_ calculation, using germination in place of toxicology data, from the Rstudio package ‘MASS’ (Venables and Ripley, 2002), was then used to calculate the time during the dormancy period at which 50 % of all viable seed had germinated (D_50_) per genotype, using the above calculation of true dormancy break.

## Results

### Evaluation of NWR Dormancy

The pattern of dormancy break over the expected 3-6 month dormancy period for NWR has not been reported previously. In this study, the percentage of germination was recorded and plotted across a 9-month period, following harvest, in order to establish a dormancy curve. The germination of Itasca-C12 and K2EF-C16 was tracked from September 2019 through May of 2020. Both seed lines tested had reached 95 % germination by January 2^nd^, approximately 4 months (17 weeks) post-harvest (Figure 2). This falls within the frequently recorded 3-6 month dormancy window for NWR. From January (~17 weeks) through the end of the study, in May (~35 weeks), the germination remained high, with means between 81-98 %. Both lines followed a similar pattern of germination through January, though K2EF-C16 continuously had higher germination through that period. The germination gap between lines began to lessen around November (~10 weeks), at which time it appears that Itasca-C12 caught up and remained comparable with K2EF-C16 for the remainder of the study. The calculated D_50_ value, or the timepoint at which 50 % of viable seed had broken dormancy, was shorter for K2EF-C16, at only 5.3 weeks, compared with that of Itasca-C12, at 6.2 weeks (Figure 2). However, both had reached their D_50_ prior to 3 months, or the expected start of dormancy break for NWR.

**Figure 2.**
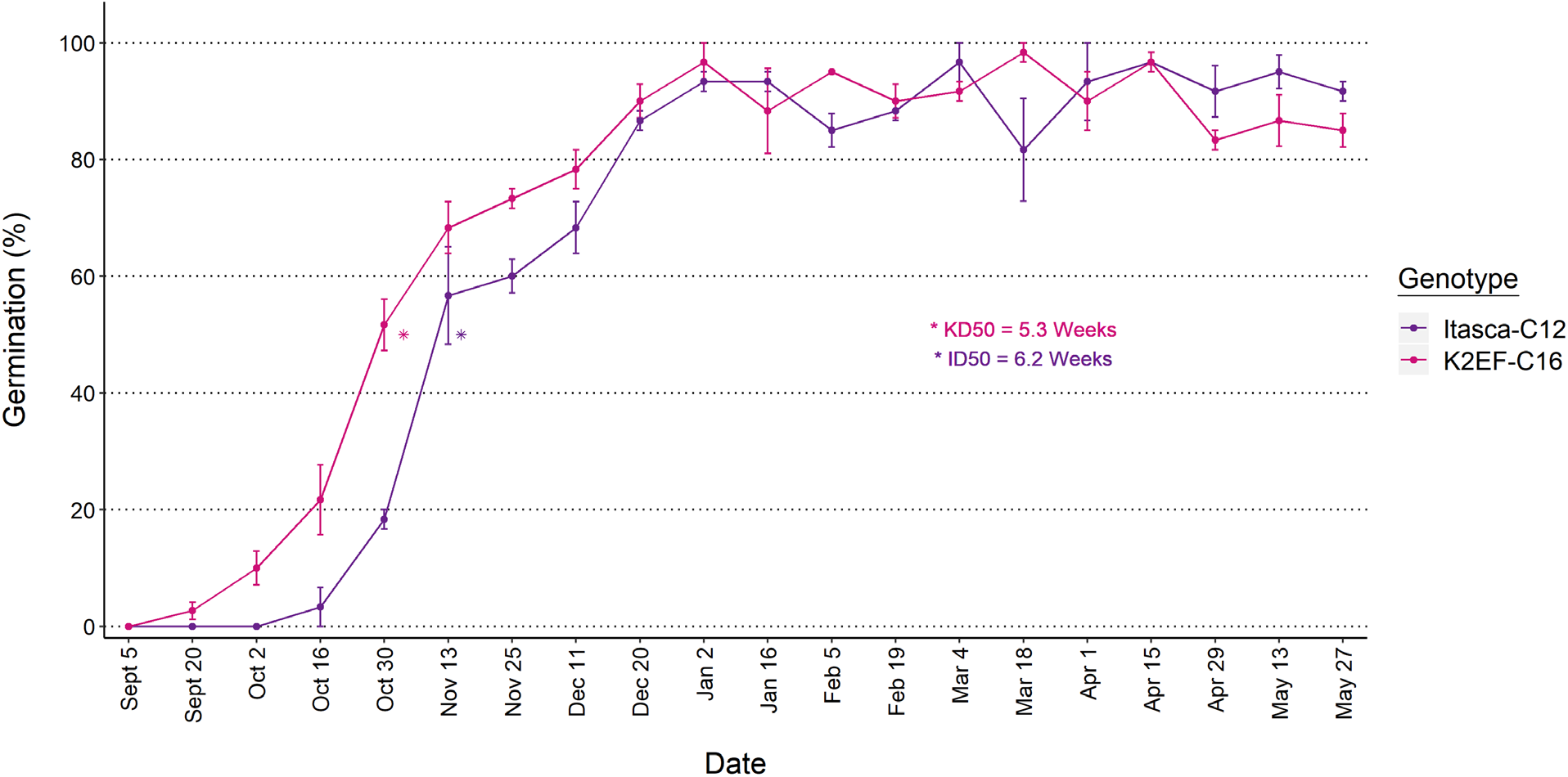
The germination of two genotypes, Itasca-C12 and K2EF-C16, of northern wild rice (NWR; *Zizania palustris* L.) beginning after harvest, on September 5^th^, 2019 and ending 9 months later, on May 27^th^, 2020. D_50_ values were a calculation of the time required for 50 % of all viable seed to germinate for each genotype.

### Dormancy Breaking Experiment – Germination and Viability

In an effort to increase the number of breeding cycles possible in a year, treatments were tested for their ability to break dormancy early in NWR. The efficacy of the combination of primary and secondary dormancy breaking treatments was assessed using results from germination tests, tetrazolium staining, and seedling and plant ratings. At 1 week post-harvest, germination across all treatment combinations was less than 15 % (Figure 3a). Germination at this point was similar to that seen in the dormancy curve (Figure 2). Although germination was low, there was a significant effect of primary treatments on the germination of seed (Table 1). Seeds treated with sulfuric acid had significantly higher germination than other primary treatments, 5.8 % compared with 0.53 % and 0 % for water and NaClO, respectively, when averaged across secondary treatments (Table 3a). There was no significant effect on germination due to secondary treatments at this point (Table 1).

**Figure 3.**
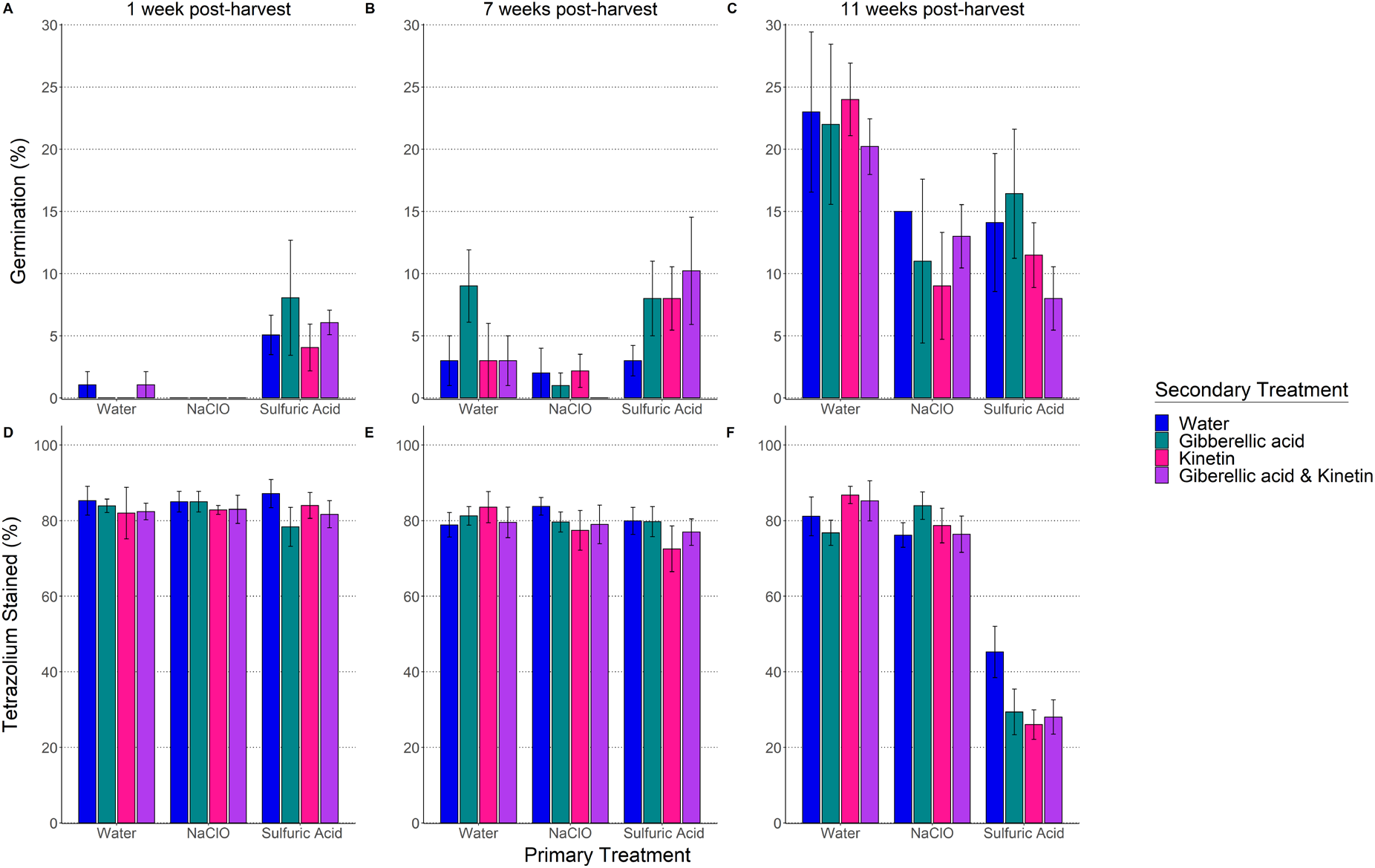
The percent of germination (A-C) and tetrazolium staining (D-F) of northern wild rice (NWR; *Zizania palustris* L.) at 1 (A and D), 7 (B and E), and 11 (C and F) weeks post-harvest, following primary and secondary dormancy treatments.

**Table 1.** Analysis of variance (ANOVA) of percent germination (%G) of treated northern wild rice (NWR; *Zizania palustris* L.) seed at 1, 7, and 11 weeks post-harvest following primary (2 hour soak in 300 ml of 2.63 % sodium hypochlorite, a 45 second wash in 200 ml of 98 % sulfuric acid, or a 45 second wash in dH_2_0) and secondary (0.1 mM gibberellic acid (GA_3_), 1 mM kinetin (K), GA_3_ and K, or dH_2_0) treatments.

| **1 week post-harvest** |  |  |  |  |  |
| --- | --- | --- | --- | --- | --- |
|  | Df | Sum sq | Mean sq | F-value | Pr(>F) |
| Rep | 4 | 28.50 | 7.12 | 0.54 | 0.710 |
| Primary Treatment | 2 | 412.77 | 206.38 | 15.52 | < 0.001 |
| Secondary Treatment | 3 | 14.69 | 4.90 | 0.37 | 0.776 |
| Primary x Secondary | 6 | 34.68 | 5.78 | 0.43 | 0.852 |
| Residuals | 44 | 584.93 | 13.29 |  |  |
| **7 weeks post-harvest** |  |  |  |  |  |
|  | Df | Sum sq | Mean sq | F-value | Pr(>F) |
| Rep | 4 | 187.74 | 46.93 | 1.76 | 0.153 |
| Primary Treatment | 2 | 361.73 | 180.87 | 6.80 | < 0.01 |
| Secondary Treatment | 3 | 83.40 | 27.80 | 1.04 | 0.382 |
| Primary x Secondary | 6 | 206.80 | 34.47 | 1.30 | 0.279 |
| Residuals | 44 | 1170.92 | 26.61 |  |  |
| **11 weeks post-harvest** |  |  |  |  |  |
|  | Df | Sum sq | Mean sq | F-value | Pr(>F) |
| Rep | 4 | 651.94 | 162.99 | 1.76 | 0.153 |
| Primary Treatment | 2 | 1349.54 | 674.77 | 7.30 | < 0.01 |
| Secondary Treatment | 3 | 119.36 | 39.79 | 0.43 | 0.732 |
| Primary x Secondary | 6 | 215.98 | 36.00 | 0.39 | 0.882 |
| Residuals | 44 | 4067.93 | 92.45 |  |  |

At 7 weeks post-harvest, the maximum germination remained below 15 % (Figure 3b), which was a little lower than the equivalent time period, Oct. 30^th^ (~8 weeks), in the dormancy curve (Figure 2). Despite the low germination, there was a significant effect of primary treatments on the germination of seed (Table 1). Sulfuric acid treated seeds had the highest average germination at 7.3 %, but the germination was not significantly different than that of water primary treated seed, at 4.5 % (Table 3b). The proximity of the germination values may be due to the interaction between water as a primary treatment and the GA_3_ secondary treatment, which had comparable germination to those of the sulfuric acid treatments, increasing the average germination of the water primary treatment (Figure 3b). Both sulfuric acid and water had higher average germination than NaClO, at only 1.3 % (Table 3b). Despite the germination peak between water and GA_3_, no significant differences in germination were attributable to secondary treatments or to the combination of primary and secondary treatments (Table 1).

At 11 weeks post-harvest, the maximum germination increased to about 30 %, which is quite a bit lower than that seen in the dormancy curve (60% for Itasca-C12 and 73% for K2EF-C16) at the equivalent time period, Nov. 25^th^ (~11.5 weeks) (Figure 3c; Figure 2). Primary treatments had a significant effect on germination (Table 1). However, the water primary treated seed now showed significantly higher germination than either sulfuric acid or NaClO treatments, 22.3 % compared to 12.5 % and 12.0 %, respectively, when averaged across secondary treatments (Table 3c). Secondary treatments still showed no significant effect on germination.

The viability of all non-germinating seed was tracked during the trials, to determine whether the lack of germination was due to dormant or dead seed. At 1 week post-harvest, there were no differences in viability between primary or secondary treatments, or the combination of the two (Table 2). For all treatments, about 80 % of the embryos stained, indicating high viability (Figure 3d). By 7 weeks post-harvest, the percent of stained seed had decreased slightly, with a range of 73-83 % (Figure 3e). Once again, no significant differences were found between treatments (Table 2). At 11 weeks post-harvest, there was a significant effect of primary treatment on the staining of seed (Table 2). While the range of staining for NaClO and water primary treated seeds remained stable, compared to the previous trial, seeds treated with sulfuric acid had significantly lower staining, with a range of just 26-45 % (Figure 3f). There was no effect of secondary treatments on the tetrazolium staining of seed.

**Table 2.** Analysis of variance (ANOVA) of percent tetrazolium-stained seed of treated northern wild rice (NWR; *Zizania palustris* L.) seed at 1, 7, and 11 weeks post-harvest, following primary (2 hour soak in 300 ml of 2.63 % sodium hypochlorite, a 45 second wash in 200 ml of 98 % sulfuric acid, or a 45 second wash in dH_2_0) and (0.1 mM gibberellic acid (GA_3_), 1 mM kinetin (K), GA_3_ and K, or dH_2_0) secondary treatments.

| **1 week post-harvest** |  |  |  |  |  |
| --- | --- | --- | --- | --- | --- |
|  | Df | Sum sq | Mean sq | F-value | Pr(>F) |
| Rep | 4 | 252.05 | 63.01 | 0.91 | 0.464 |
| Primary Treatment | 2 | 13.69 | 6.85 | 0.10 | 0.906 |
| Secondary Treatment | 3 | 120.09 | 40.03 | 0.58 | 0.631 |
| Primary x Secondary | 6 | 140.68 | 23.45 | 0.34 | 0.912 |
| Residuals | 44 | 3034.34 | 68.96 |  |  |
| **7 weeks post-harvest** |  |  |  |  |  |
|  | Df | Sum sq | Mean sq | F-value | Pr(>F) |
| Rep | 4 | 584.43 | 146.11 | 1.96 | 0.118 |
| Primary Treatment | 2 | 134.36 | 67.18 | 0.90 | 0.414 |
| Secondary Treatment | 3 | 89.82 | 29.94 | 0.40 | 0.753 |
| Primary x Secondary | 6 | 264.19 | 44.03 | 0.59 | 0.737 |
| Residuals | 44 | 3284.63 | 74.65 |  |  |
| **11 weeks post-harvest** |  |  |  |  |  |
|  | Df | Sum sq | Mean sq | F-value | Pr(>F) |
| Rep | 4 | 108.20 | 27.05 | 0.24 | 0.917 |
| Primary Treatment | 2 | 31468.27 | 15734.14 | 136.98 | < 0.001 |
| Secondary Treatment | 3 | 186.59 | 62.20 | 0.54 | 0.656 |
| Primary x Secondary | 6 | 1480.82 | 246.80 | 2.15 | 0.067 |
| Residuals | 44 | 5054.07 | 114.87 |  |  |

**Table 3.** Tukey’s LSD, with a Bonferroni adjustment to p-values, for all significant effects of dormancy breaking treatments on the percent of germination (%G) or tetrazolium (Tz) staining of seed from northern wild rice (NWR; *Zizania palustris* L.), at 1, 7, and 11 weeks post-harvest.

| **A. 1 week post-harvest** |  |  |
| --- | --- | --- |
| *Primary Treatments* | *Average Germination (%)* | *Groups* |
| Water | 0.53 | b |
| Sulfuric Acid | 5.81 | a |
| NaClO | 0.00 | b |
| **B. 7 weeks post-harvest** |  |  |
| *Primary Treatments* | *Average Germination (%)* | *Groups* |
| Water | 4.50 | ab |
| Sulfuric Acid | 7.31 | ab |
| NaClO | 1.30 | b |
| **C. 11 weeks post-harvest** |  |  |
| *Primary Treatments* | *Average Germination (%)* | *Groups* |
| Water | 22.3 | a |
| Sulfuric Acid | 12.50 | b |
| NaClO | 12.00 | b |
| *Primary Treatments* | *Average Tz Staining (%)* | *Groups* |
| Water | 82.48 | a |
| Sulfuric Acid | 32.17 | b |
| NaClO | 78.81 | a |

### Dormancy Breaking Experiment – Seedling and Plant Health

For the purpose of breeding, it is not enough to break dormancy early, the resulting plants must also remain viable thereafter. Therefore, seedling and plant health was assessed for all seed that germinated during the trials. At 1 week post-harvest, only 23 seeds germinated, resulting in a small data set for this time point. Of the germinated seeds, the treatment combination resulting in the longest seedling was a water primary treatment and a secondary treatment with GA_3_ and Kinetin (Figure 4a). However, there was only one seed to measure for this treatment, making this data difficult to interpret. Many of the seeds that germinated at this time, were those treated with a sulfuric acid primary treatment. Of the germinated seeds treated with sulfuric acid, the secondary treatment with the lowest mean seedling length, 16.91 cm, was kinetin, while the highest, at 26.08 cm, was GA_3_ and kinetin (Table S1). The only plant during this trial that had a length above 0, by the time plant ratings were collected, was that treated with sulfuric acid, followed by GA_3_. It had a plant length of 451.11 cm and a plant health rating of 3 (Figure 4d; 5g). All remaining plants had died and had a length of 0 cm and a health rating of 1.

**Figure 4.**
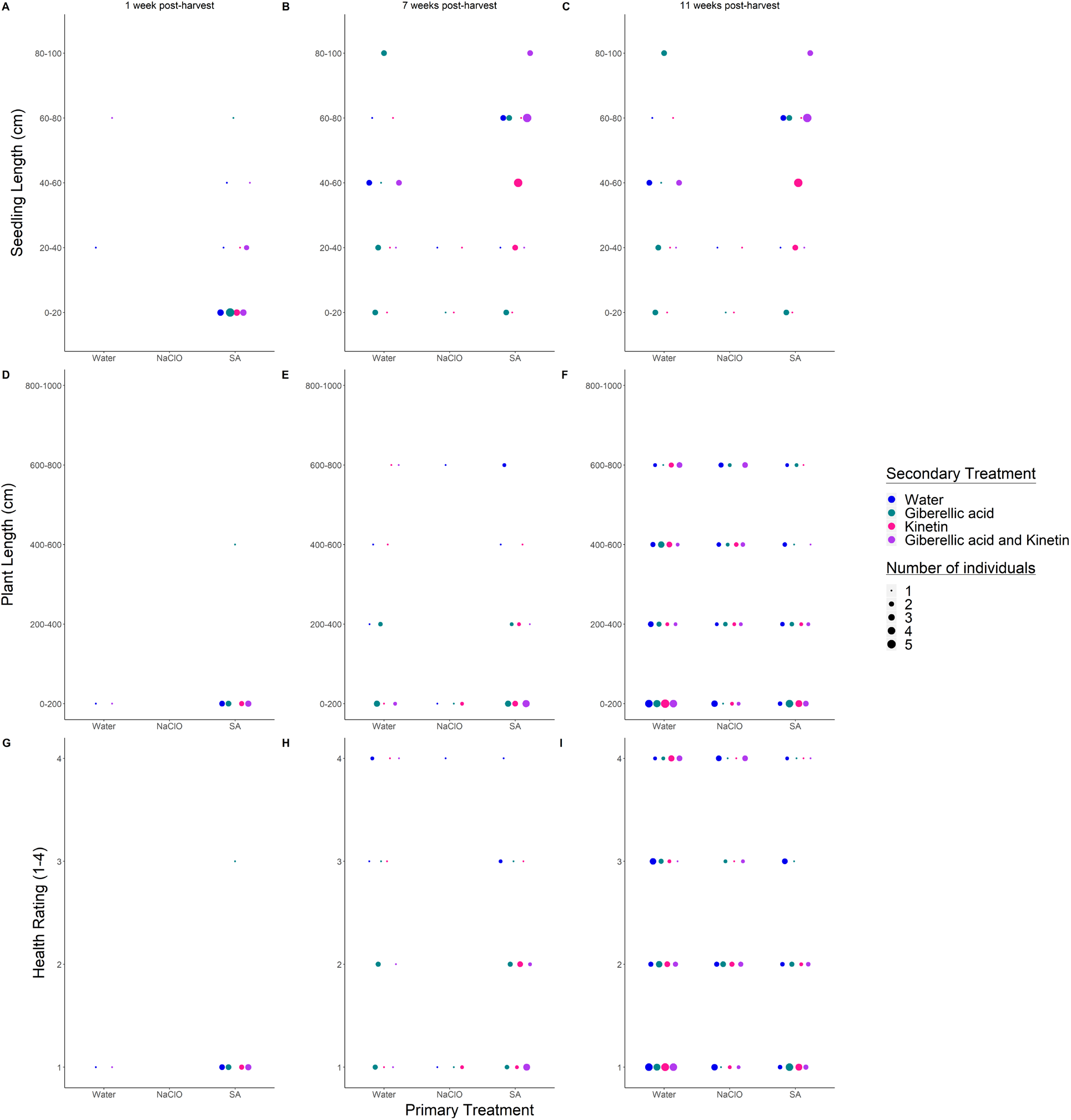
The seedling length (A-C), plant length (D-F), and plant health ratings (G-I) of northern wild rice (NWR; *Zizania palustris* L.) at 1 (A, D, and G), 7 (B, E and H), and 11 (C, F, and I) weeks post-harvest following primary and secondary dormancy treatments.

At 7 weeks post-harvest, 52 seeds germinated, with representation from all primary treatments (Figure 3a). The seedlings with the highest mean length, 94.61 cm (n=8), were treated with SA and GA_3_ and the lowest, at 18.72 cm, was treated with NaClO and GA_3_ (n=1) (Supplementary Table 1). Although some seedlings treated with water and GA_3_ did have lengths matching the highest overall mean length, others were much shorter, resulting in a lower mean for this treatment combination (Figure 4b; Supplementary Table 1). Overall, seeds treated with NaClO resulted in short seedlings (≤ 40 cm), relative to other primary treatments. Seedling length did not appear to predict plant length, as seedlings with the highest mean seedling length, SA with GA_3_, resulted in one of the shortest mean plant lengths (Supplementary Table 1). The highest mean plant length, 631.80 cm, resulted from seeds treated with SA, then water, closely followed by those treated with water, then water, at 563.23 cm (Supplementary Table 1). However, ~63 % of seedlings from this trial resulted in short plants (≤ 200) and 75 % had plant health ratings of 1 or 2 (Figure 4e; 5h).

By 11 weeks post-harvest 182 seeds had germinated, a marked increase from either of the two previous trials. As these seeds were distributed across all primary and secondary treatments, it appeared that seed may have been naturally beginning the process of seed dormancy break. However, as with the previous trial, the seedlings with the highest mean length, 93.96 cm, were treated with SA, then GA_3_, although the lowest were treated with NaClO, then water (Supplementary Table 1). In the previous trial, seeds treated with NaClO had a higher rate of plant death, relative to other primary treatments, but by the 11 week post-harvest trial, seeds treated with NaClO actually had a higher mean plant length than either other primary treatment, when averaged across all secondary treatments (Supplementary Table 1). Interestingly, the longest mean seedling lengths, still resulted in one of the shortest mean plant lengths (Supplementary Table 1). However, in general seedling length was not a good predictor of plant length. During this trial ~ 44 % of seedlings resulted in short plants and ~69 % of plants had a plant health rating of 1 or 2 (Figure 4f; 5i).

## Discussion

### Evaluation of NWR Dormancy

An understanding of when dormancy break begins, peaks, and ends will help researchers to form a more complete picture of the process of dormancy break in NWR. The dormancy curve developed in this study showed that from January (~17 weeks post-harvest) through the end of the study, in May (~35 weeks post-harvest), the germination remained high, indicating that no significant loss in viability had taken place. However, because this study only covered about 9 months post-harvest, it was likely too short to demonstrate the drop in viability that is commonly recorded for NWR (Kovach and Bradford, 1992a; Berjak and Pammenter, 2008; McGilp *et al*., 2020). Additionally, the D_50_ for K2EF-C16 was shorter than that of Itasca-C12. This is consistent with previous anecdotal observations within the breeding program, in which K2EF-C16 has broken dormancy early, relative to Itasca-C12, which falls closer to the average dormancy break of breeding germplasm. However, it is interesting that the D_50_ occurred at less than 2 months following harvest, given the expected 3-6 month dormancy period for NWR (Simpson, 1966; Cardwell *et al*., 1978; Kovach and Bradford, 1992a). While departing from the expected range, intraspecific and intrapopulation variation in the dormancy and germination of seed is also found in many other species, including *Linum perenne* L, *Spergula arvensis*, *Sinapis arvensis*, and *Viminaria juncea* (Meyer and Kitchen, 1994; Andersson and Milberg, 1998; Jones and Nielson, 1999; Baskin and Baskin, 2004; Lacerda *et al*., 2004; Liyanage and Ooi, 2015). Given the heterogeneous and heterozygous nature of open-pollinated breeding populations, these types of variation are likely to exist in NWR.

### Dormancy Breaking Experiment – Germination and Viability

Seed dormancy and germination are traits controlled by both genetics and a variety of environmental factors (Koornneef *et al*., 2002; Skubacz, 2017). Many genes are involved in the initiation, maintenance, and decline of seed dormancy, which are regulated by the presence or absence of hormones and certain environmental cues including but not limited to: high or low temperatures, oxygen or nitrate availability, and the time for embryo growth and expansion (Baskin and Baskin, 2004; Finch-Savage and Leubner-Metzger, 2006; Finkelstein *et al*., 2008; Duermeyer *et al*., 2018). Although there are various types of dormancy, at a basic level, the regulation of dormancy involves the interplay of the growth potential of the embryo and its physical or chemical restraint by the surrounding seed tissues (Kucera *et al*., 2007; Baskin and Baskin, 2014a).

Unique breeding challenges exist for NWR due to the seeds’ intermediate recalcitrance and the presence of seed dormancy. Previous studies have evaluated these traits in order to improve the understanding of NWR seed physiology and thereby increase breeding efficiency (Cardwell *et al*., 1978; Oelke and Albrecht, 1978; Albrecht *et al*., 1979; Kovach and Bradford, 1992a; b; McGilp *et al*., 2020). However, due to the relatively recent cultivation of NWR compared to other crops, and the unique combination of recalcitrance and dormancy, the mechanisms underlying dormancy in NWR are not well understood. This study sought to identify effective and efficient methods of early dormancy break in NWR for the purpose of breeding, the results of which could also hint at the types of dormancy regulation present in the seed.

Germination, viability, and seedling and plant health were tracked starting at 1 week post-harvest and continuing through 11 weeks post-harvest, to determine the efficacy of primary and secondary treatments for early dormancy break in NWR. Across all timepoints germination was low, never reaching above 30 %, regardless of treatment, although tetrazolium tests indicated that this was not a matter of seed viability. Such low germination would not provide adequate plants for breeding purposes. However, some differences were seen between primary treatments, though none were identified between secondary treatments at any time point.

At 1 week post-harvest, the sulfuric acid treated seed had significantly higher germination than NaClO or water, when averaged across secondary treatments. Sulfuric acid is effective in breaking seed dormancy in a number of species with physical dormancy including field bindweed (*Convolvulus arvensis* L.), *Medicago* spp., and *Trifolium* spp., among others (Balouchi and Sanavy, 2006; Kimura and A, 2012; Baskin and Baskin, 2014b; Xiong *et al*., 2018). No NaClO treated seed germinated at this timepoint, perhaps indicating that NaClO was not sufficient to break through or weaken the seed coats at their thickest point during the dormancy period to allow imbibition to take place. While previous results have indicated that treatment of freshly harvested NWR seed with NaClO was effective for germination, these studies have used dehulled and scarified seed, effectively providing a double scarification treatment (Oelke and Albrecht, 1980). Additionally, while NaClO is effective for dormancy break in some species, in others it can have a negative effect on germination, and in some it can have no effect whatsoever, even within the same plant families (Rosbakh *et al*., 2019). By 7 weeks post-harvest sulfuric acid treated seed did not have significantly higher germination than the primary water treated seed but both still had higher germination than NaClO treated seed. At 11 weeks post-harvest the water treated seed had significantly higher germination than either of the other primary treatments. This may indicate that by this time point the embryo covering layers were undergoing natural weakening processes, perhaps triggered by sufficient cold stratification, allowing for further growth of the embryo, as occurs in seeds with physiological dormancy (Bewley *et al*., 2013; Baskin and Baskin, 2014a). As this was close to 3 months post-harvest, it falls approximately within the expected timeframe for dormancy break in NWR (Simpson, 1966; Cardwell *et al*., 1978; Oelke and Albrecht, 1980).

The viability of all seed, estimated by the tetrazolium staining of embryos, remained high (~80%) through the 7 week post-harvest time point, regardless of treatment. As this is very shortly after harvest, high viability was expected. However, by 11 weeks post-harvest, the primary treatment had a significant effect on the viability of seed, with water treated seed displaying the highest viability, when averaged across secondary treatments. Seeds treated with NaClO had similar levels of viability to water treated seed, though lower on average. Water and NaClO treated seed retained levels of viability near that of the previous timepoints, whereas SA treated seeds had significantly lower viability. This likely indicates that SA had a deleterious effect on the viability of seed by the 11 week timepoint, most likely due to the SA reaching and damaging the embryo. Additionally, seed treated with NaClO had lower germination than that treated with water but the seed showed no real loss in viability, perhaps indicating that NaClO was not truly an effective treatment, so much as a neutral one. Previous research has also shown only a modest effect of NaClO on the germination of NWR seed, even when combined with mechanical scarification (Oelke and Albrecht, 1980).

While seeds were viable and able to germinate this early in the dormancy period, they did not result in viable seedlings. Previous NWR seed research has also found that while dormancy can be broken shortly after harvest by the removal of structures covering the embryo, the resulting seedlings are weak (Oelke and Albrecht, 1980). Little research has been devoted to excising embryos of NWR to assess the growth of seedlings. One study found that when embryos of NWR seed were excised, they could germinate normally, perhaps indicating that the physiological dormancy is at a non-deep level (LaRue and Avery, 1983). However, it is unknown at what point in development these seeds were harvested or whether the germinated seeds would have resulted in viable plants. Additionally, the results have not been replicated. The inability of an excised embryo to produce normal seedlings is one of the factors used to differentiate between intermediate and deep physiological dormancy, though it is often difficult to distinguish between the levels of physiological dormancy (Baskin and Baskin, 2004, 2014a).

Based upon the results of this study, we propose that NWR exhibits intermediate or deep physiological dormancy, requiring sufficient cold stratification in order to germinate and produce viable seedlings (Baskin and Baskin, 2014a). Previous studies have concluded that NWR must have physical dormancy, due to the appearance of an impermeable seed coat and the observation that removal of certain portions of the seed coat can induce germination (Simpson, 1966; Cardwell *et al*., 1978; Oelke and Albrecht, 1978). However, the actual imbibition of NWR seed has not been directly measured (Dalziell *et al*., 2019), nor has the structure of the NWR pericarp, in order to determine with confidence that it is truly impermeable. Additionally, a review of nearly 900 tree and shrub species found that while about 14 % of desiccation sensitive seeds displayed physiological dormancy, only 1.4 % of the species had physical dormancy, resulting from an impermeable seed coat (Tweddle *et al*., 2003; Jaganathan, 2021). While that study found that most desiccation sensitive seed displayed morphological or morphophysiological dormancy, we believe that the embryo of dormant NWR seed is both differentiated and developed, based upon the pattern of embryo staining when tetrazolium is applied shortly after harvest. However, a more complete study of embryo development is needed to confirm these anecdotal observations. Therefore, while previous studies have stated otherwise, it is possible that the constraint of the embryo covering layers is not due to impermeability but rather to the mechanical constraint imposed by the seed coat or the low growth potential of the embryo at harvest, which is alleviated over time in cold stratification. Previous research has also demonstrated the need for cold stratification for dormancy break in NWR (Kovach and Bradford, 1992a; McGilp *et al*., 2020). In *Najas marina* and *Trapella sinensis*, two annual, aquatic plants with seed dormancy, cold stratification was also required for the termination of dormancy, leading the authors to suggest that their seed may by physiologically dormant (Handley and Davy, 2005; Kato and Kadono, 2011). While there are species whose seeds have combinational dormancy, or both physiological and physical dormancy, it is rare and unlikely to occur when deep dormancy is present. In addition, combinational dormancy within desiccation sensitive seed was entirely absent in a dataset of 900 tree and brush species (Tweddle *et al*., 2003; Baskin and Baskin, 2004). Regardless of the cause of the restriction to dormancy imposed by the seed coat, researchers have noted that damage to the pericarp is helpful but not sufficient for successful dormancy break and subsequent survival of seedlings in NWR (Simpson, 1966; Grombacher *et al*., 1997).

In this experiment, secondary hormone treatments showed no significant effect on the germination of dormant seed, possibly indicating that hormonal signaling is not the limiting factor for dormancy break in NWR. A lack of dormancy break in response to GA_3_ can indicate that seed has deep or intermediate PD, as has been found in some Forbes species as well as *Rosa multibracteata* (Baskin and Baskin, 2004; Hoyle *et al*., 2008; Zhou *et al*., 2009). It is also possible that different hormones are involved in the dormancy process of NWR, as ethylene, brassinosteroids, and nitrate are also known to be involved in the cessation of dormancy in other species, or that the hormones used in this study were applied at ineffective rates (Kucera *et al*., 2007; Baskin and Baskin, 2014a; Toorop, 2015). While there were some significant interactions between primary and secondary treatments in relation to seedling and plant health, they were not consistent across time, making it difficult to draw any broad conclusions about their efficacy. Previous studies have found mixed results on the efficacy of the application of GA_3_, with or without disruption of the seed coat, for the induction of dormancy break in NWR (Simpson, 1966; Oelke and Albrecht, 1980). However, ABA has been found in high concentrations in the hulls, pericarps, and embryos of dormant NWR seed and the application of hulls and pericarps from freshly harvested seed to non-dormant seed has resulted in the inhibition of germination (Cardwell *et al*., 1978; Albrecht *et al*., 1979). This indicates that ABA is involved in the induction of dormancy in NWR but the connection between GA_3_ and other tested hormones for the efficacious reduction of dormancy of large NWR seed lots is less clear. As the effect of GA_3_ application on dormancy break is one of the differentiating factors between non-deep and deep PD (Baskin and Baskin, 2004), the inconclusiveness of these tests further complicates the ability to characterize dormancy in NWR. Ultimately, sufficient time under cold stratification conditions has been shown to be the most effective strategy for the production of sufficient healthy NWR plants for the purpose of breeding (Grombacher *et al*., 1997).

Therefore, early forced germination of the types tested here and previously for NWR are unlikely to result in a sufficient number of healthy NWR plants for the purpose of breeding. Transcriptomic analysis over the course of NWR dormancy, as has been conducted in *Chenopodium quinoa* and *Panax quinquefolius* (Qi *et al*., 2015; Wu *et al*., 2020), is one tool that should be explored in the future, to better understand the dormancy process in NWR and to develop more efficacious strategies for increasing the speed of breeding. Additionally, characterization of the imbibition of dormant and non-dormant seed and the physical structure of the embryo and surrounding tissues will further illuminate the presence or absence of physical dormancy. Such strategies will elucidate active pathways and physical barriers throughout the course of dormancy, ultimately informing future breeding strategies.

## Conclusion

A dormancy curve for NWR has been developed across 9 months following harvest. This study gave insight into the type of dormancy present in NWR. While previous research has concluded that NWR has physical dormancy, evidence from this and previous studies point to a physiological source of dormancy. Due to the focus on treatments aimed at physical dormancy, there has been little headway made in the production of healthy plants through early dormancy break, for breeding of NWR. Future studies assessing the imbibition, physical structure, and expression levels of dormant and non-dormant NWR seed will allow for improved methodology development, leading to more efficient breeding strategies.

---

**Supplementary Table 1.** The number of individuals tested, mean seedling length, and mean plant length of northern wild rice (NWR; *Zizania palustris* L.) at 1, 7, or 11 weeks post-harvest and following primary and secondary dormancy treatments.

| Trial | Primary Treatment | Secondary Treatment | Number of individuals | Mean seedling length (cm) | Mean plant length (cm) |
| --- | --- | --- | --- | --- | --- |
| 1 week post-harvest | Water | Water | 1 | 21.24 | 0.00 |
|  | Water | GA and Kinetin | 1 | 73.76 | 0.00 |
|  | Sulfuric Acid | Water | 5 | 20.42 | 0.00 |
|  | Sulfuric Acid | GA | 6 | 22.17 | 75.19 |
|  | Sulfuric Acid | Kinetin | 4 | 16.91 | 0.00 |
|  | Sulfuric Acid | GA and Kinetin | 6 | 26.08 | 0.00 |
| 7 weeks post-harvest | Water | Water | 3 | 57.69 | 563.23 |
|  | Water | GA | 9 | 65.65 | 132.42 |
|  | Water | Kinetin | 3 | 38.63 | 372.33 |
|  | Water | GA and Kinetin | 3 | 50.42 | 286.43 |
|  | NaClO | Water | 2 | 71.63 | 354.65 |
|  | NaClO | GA | 1 | 18.72 | 0.00 |
|  | NaClO | Kinetin | 2 | 29.33 | 0.00 |
|  | Sulfuric Acid | Water | 3 | 58.76 | 631.80 |
|  | Sulfuric Acid | GA | 8 | 94.61 | 119.51 |
|  | Sulfuric Acid | Kinetin | 8 | 41.16 | 180.60 |
|  | Sulfuric Acid | GA and Kinetin | 10 | 58.83 | 32.41 |
| 11 weeks post-harvest | Water | Water | 23 | 37.74 | 283.61 |
|  | Water | GA | 22 | 45.06 | 260.00 |
|  | Water | Kinetin | 24 | 35.08 | 248.86 |
|  | Water | GA and Kinetin | 20 | 47.04 | 298.77 |
|  | NaClO | Water | 16 | 29.63 | 313.25 |
|  | NaClO | GA | 11 | 67.75 | 489.32 |
|  | NaClO | Kinetin | 9 | 32.26 | 372.08 |
|  | NaClO | GA and Kinetin | 13 | 47.85 | 496.04 |
|  | Sulfuric Acid | Water | 14 | 53.67 | 430.55 |
|  | Sulfuric Acid | GA | 17 | 93.96 | 164.00 |
|  | Sulfuric Acid | Kinetin | 11 | 34.48 | 129.61 |
|  | Sulfuric Acid | GA and Kinetin | 8 | 69.07 | 211.74 |

## References

Albrecht, K. A., Oelke, E. A. and Brenner, M. L. (1979) Abscisic acid levels in the grain of wild rice. Crop Science 19, 671–676. doi:10.2135/cropsci1979.0011183X001900050032x.

Andersson, L. and Milberg, P. (1998) Variation in seed dormancy among mother plants, populations and years of seed collection. Seed Science Research 8, 29–38. doi:DOI: 10.1017/S0960258500003883.

Balouchi, H. R. and Sanavy, S. A. M. M. (2006) Effect of gibberellic acid, prechilling, sulfuric acid and potassium nitrate on seed germination and dormancy of annual medics. Pakistan Journal of Biological Sciences 9, 2875–2880. doi:10.3923/pjbs.2006.2875.2880.

Baskin, J. M. and Baskin, C. C. (2004) A classification system for seed dormancy. Seed Science Research 14, 1–16. doi:DOI: 10.1079/SSR2003150.

Baskin, C. C. and Baskin, J. M. (2014a) Chapter 3 - Types of seeds and kinds of seed dormancy, pp. 37–77 in Baskin, C. C. and Baskin, J. M. (Eds.) Seeds: Ecology, biogeography, and evolution of dormancy and germination. San Diego, Academic Press doi:https://doi.org/10.1016/B978-0-12-416677-6.00003-2.

Baskin, C. C. and Baskin, J. M. (2014b) Chapter 6 - Germination ecology of seeds with physical dormancy, pp. 145–185 in Baskin, C. C. and Baskin, J. M. (Eds.) Seeds: Ecology, biogeography, and evolution of dormancy and germination. San Diego, Academic Press doi:https://doi.org/10.1016/B978-0-12-416677-6.00006-8.

Berjak, P. and Pammenter, N. W. (2008) From Avicennia to Zizania: Seed recalcitrance in perspective. Annals of Botany 101, 213–228. doi:10.1093/aob/mcm168.

Bewley, J. D., Bradford, K. J., Hilhorst, H. W. M. and Nonogaki, H. (2013) Dormancy and the control of germination, pp. 247–297 in Bewley, J. D., Bradford, K. J., Hilhorst, H. W. M., and Nonogaki, H. (Eds.) Seeds: Physiology of Development, Germination and Dormancy. New York, NY, Springer New York doi:10.1007/978-1-4614-4693-4_6.

Biesboer, D. D. (2019) The ecology and conservation of wild rice, Zizania palustris l., in North America. Acta Limnologica Brasiliensia 31, e102.

Cardwell, V. B., Oelke, E. A. and Elliott, W. A. (1978) Seed dormancy mechanisms in wild rice (Zizania aquatica). Agronomy 70, 481–484.

Cohn, M. A. and Butera, D. L. (1982) Seed dormancy in red rice (Oryza sativa). II. Response to cytokinins. Weed Science 30, 200–205.

Dalziell, E. L., Baskin, C. C., Baskin, J. M., Young, R. E., Dixon, K. W. and Merritt, D. J. (2019) Morphophysiological dormancy in the basal angiosperm order Nymphaeales. Annals of botany 123, 95–106. doi:10.1093/aob/mcy142.

Dickie, J. B. and Pritchard, H. W. (2002) Systematic and evolutionary aspects of desiccation tolerance in seeds, pp. 239–260 in Black, M. and Pritchard, H. W. (Eds.) Dessication and Survival in Plants: Drying Without Dying. CABI.

Duermeyer, L., Khodapanahi, E., Yan, D., Krapp, A., Rothstein, S. J. and Nambara, E. (2018) Regulation of seed dormancy and germination by nitrate. Seed Science Research 28, 150–157. doi:https://doi.org/10.1017/S096025851800020X.

Finch-Savage, W. E. and Leubner-Metzger, G. (2006) Seed dormancy and the control of germination. The New Phytologist 171, 501–523.

Finkelstein, R., Reeves, W., Ariizumi, T. and Steber, C. (2008) Molecular aspects of seed dormancy. Annual Review of Plant Biology 59, 387–415. doi:10.1146/annurev.arplant.59.032607.092740.

Grombacher, A. W., Porter, R. A. and Everett, L. A. (1997) Breeding wild rice. Plant Breeding Reviews 14 237–265. doi:10.1002/9780470650073.ch8.

Handley, R. J. and Davy, A. J. (2005) Temperature effects on seed maturity and dormancy cycles in an aquatic annual, Najas marina, at the edge of its range. Journal of Ecology 93, 1185–1193.

Hilhorst, H. W. M. (1995) A critical update on seed dormancy. I. Primary dormancy. Seed Science Research 5, 61–73. doi:DOI: 10.1017/S0960258500002634.

Hoyle, G. L., Steadman, K. J., Daws, M. I. and Adkins, S. W. (2008) Physiological dormancy in forbs native to south–west Queensland: Diagnosis and classification. South African Journal of Botany 74, 208–213. doi:https://doi.org/10.1016/j.sajb.2007.11.005.

Huseby, J. T. (1997) Use of cultivated wild rice paddies and associated habitats by migrating and breeding waterfowl in northwest Minnesota. http://login.ezproxy.lib.umn.edu/login? url=https://www.proquest.com/dissertations-theses/use-cultivated-wild-rice-paddies-associated/docview/304378734/se-2?accountid=14586.

Jaganathan, G. K. (2021) Ecological insights into the coexistence of dormancy and desiccation-sensitivity in Arecaceae species. Annals of Forest Science 78, 10. doi:10.1007/s13595-021-01032-9.

Jones, T. A. and Nielson, D. C. (1999) Intrapopulation genetic variation for seed dormancy in indian ricegrass. Journal of Range Management 52, 646–650. doi:10.2307/4003636.

Kato, R. and Kadono, Y. (2011) Seed germination traits of Trapella sinensis (Trapellaceae), an endangered aquatic plant in Japan: Conservation implications. Aquatic Botany 95, 258–261. doi:https://doi.org/10.1016/j.aquabot.2011.08.002.

Kimura, E. and A, I. M. (2012) Seed scarification methods and their use in forage legumes. Research Journal of Seed Science 5, 38–50. doi:10.3923/rjss.2012.38.50.

Koornneef, M., Bentsink, L. and Hilhorst, H. (2002) Seed dormancy and germination. Current Opinion in Plant Biology 5, 33–36. doi:https://doi.org/10.1016/S1369-5266(01)00219-9.

Kovach, D. A. and Bradford, K. J. (1992a) Temperature dependence of viability and dormancy of Zizania palustris var. interior seeds stored at high moisture contents. Annals of Botany 69, 297–301.

Kovach, D. A. and Bradford, K. J. (1992b) Imbibitional damage and desiccation tolerance of wild rice *(Zizania palustris)* seeds. Journal of Experimental Botany 43, 747–757. doi:10.1093/jxb/43.6.747.

Kucera, B., Cohn, M. A. and Leubner-Metzger, G. (2007) Plant hormone interactions during seed dormancy release and germination. Seed science research 15, 281–307. doi:10.1079/SSR2005218.

Lacerda, D. R., Lemos Filho, J. P., Goulart, M. F., Ribeiro, R. A. and Lovato, M. B. (2004) Seed-dormancy variation in natural populations of two tropical leguminous tree species: Senna multijuga (Caesalpinoideae) and Plathymenia reticulata (Mimosoideae). Seed Science Research 14, 127–135. doi:DOI: 10.1079/SSR2004162.

LaRue, C. D. and Avery, G. S. (1983) The development of the embryo of Zizania aquatica in the seed and in artificial culture. Bulletin of the Torrey Botanical Club 65, 11–21.

Lata, S., Sharma, G., Garg, S. and Joshi, M. (2018) Effect of different chemical treatments on germination of strawberry seeds. International Journal of Current Microbiology and Apllied Sciences 7, 1270–1724. doi:https://doi.org/10.20546/ijcmas.2018.703.150.

Li, X., Baskin, J. M. and Baskin, C. C. (1999) Anatomy of two mechanisms of breaking physical dormancy by experimental treatments in seeds of two North American Rhus species (Anacardiaceae). American Journal of Botany 86, 1505–1511. doi:https://doi.org/10.2307/2656788.

Liyanage, G. S. and Ooi, M. K. J. (2015) Intra-population level variation in thresholds for physical dormancy-breaking temperature. Annals of botany 116, 123–131. doi:10.1093/aob/mcv069.

McGilp, L., Duquette, J., Braaten, D., Kimball, J. and Porter, R. (2020) Investigation of variable storage conditions for cultivated northern wild rice and their effects on seed viability and dormancy. Seed Science Research. doi:10.1017/S0960258520000033.

Mendiburu, F. de (2019) Agricolae: Statistical procedures for agricultural research.

Meyer, S. E. and Kitchen, S. G. (1994) Life history variation in blue flax (Linum perenne: Linaceae): Seed germination phenology. American Journal of Botany 81, 528–535. doi:10.2307/2445726.

Miyoshi, K. and Sato, T. (1997) The effects of kinetin and gibberellin on the germination of dehusked seeds of indica and japonica rice (Oryza sativa L.) under anaerobic and aerobic conditions. Annals of Botany 80, 479–483. doi:https://doi.org/10.1006/anbo.1997.0470.

Oelke, E. A. (1982) Wild rice production in Minnesota. St. Paul, Minn., University of Minnesota, Agricultural Extension Service.

Oelke, E. A. and Albrecht, K. A. (1978) Mechanical scarification of dormant wild rice seed. Agronomy Journal 70, 691–694.

Oelke, E. A. and Albrecht, K. A. (1980) Influence of chemical seed treatments on germination of dormant wild rice seeds. Crop Science 20, 595–598.

Oelke, E. A. and Porter, R. A. (2016) Wildrice, Zizania: Overview, pp. 130–139 in Corke, H., Faubion, J., W, W. C., and Seetharaman, K. (Eds.) Encyclopedia of Food Grains. Kidlington, Oxford, UK, Academic Press.

Peters, J. ed. (2000) Tetrazolium testing handbook, in Contributions to the Handbook on seed testing; no. 29. Lincoln, NE, Association of Official Seed Analysts: Tetrazolium testing committee.

Probert, R. J. and Brierley, E. R. (1989) Desiccation intolerance in seeds of Zizania palustris is not related to developmental age or the duration of post-harvest storage. Annals of Botany 64, 669–674.

Purohit, S., Nandi, S. K., Palni, L. M. S., Giri, L. and Bhatt, A. (2015) Effect of sulfuric acid treatment on breaking of seed dormancy and subsequent seedling establishment in Zanthoxylum armatum dc: an endangered medicinal plant of the himalayan region. National Academy Science Letters 38, 301–304. doi:10.1007/s40009-015-0349-5.

Qi, J., Sun, P., Liao, D., Sun, T., Zhu, J. and Li, X. (2015) Transcriptomic analysis of American ginseng seeds during the dormancy release process by RNA-Seq. PLoS ONE 10, 1–17. doi:10.1371/journal.pone.0118558.

Rosbakh, S., Hülsmann, L., Weinberger, I., Bleicher, M. and Poschlod, P. (2019) Bleaching and cold stratification can break dormancy and improve seed germination in Cyperaceae. Aquatic Botany 158, 103128. doi:https://doi.org/10.1016/j.aquabot.2019.103128.

Rstudio-team (2019) Rstudio: Integrated development for R.

Simpson, G. M. (1966) A study of germination in the seed of wild rice (Zizania aquatica). Canadian Journal of Botany 44, 1–9.

Skubacz, A. (2017) Seed dormancy : The complex process regulated by abscisic acid, gibberellins, and other phytohormones that makes seed germination work, in Mohamed El-Esawi (Ed.) Phytohormones Signaling Mechanisms and Crosstalk in Plant Development and Stress Responses. IntechOpen doi:10.5772/intechopen.68735.

Steeves, T. A. (1952) Wild rice: Indian food and a modern delicacy. Economic Botany 6, 107–142.

Toorop, P. E. (2015) Nitrate controls testa rupture and water content during release of physiological dormancy in seeds of Sisymbrium officinale (L.) Scop. Seed Science Research 25, 138–146. doi:DOI: 10.1017/S0960258514000397.

Tweddle, J. C., Dickie, J. B., Baskin, C. C. and Baskin, J. M. (2003) Ecological aspects of seed desiccation sensitivity. Journal of Ecology 91, 294–304.

Venables, W. N. and Ripley, B. D. (2002) Modern applied statistics with S. Fourth. New York, Springer.

Vieira, A. R., Vieira, M. das G. G. C., Fraga, A. C., Oliveira, J. A. and Santos, C. D. dos (2002) Action of gibberellic acid (GA3) on dormancy and activity of alpha-amylase in rice seeds. Revista brasileira de sementes 24, 43–48. doi:10.1590/S0101-31222002000100008.

Wikham, H. (2016) ggplot2-Elegant graphics for data analysis. New York, Springer-Verlag.

Woods, D. L. and Gutek, L. H. (1974) Germinating wild rice. Canadian Journal of Plant Science 54, 423–424. doi:10.4141/cjps74-064.

Wu, Q., Bai, X., Wu, X., Xiang, D., Wan, Y., Luo, Y., Shi, X., Li, Q., Zhao, J., Qin, P., Yang, X. and Zhao, G. (2020) Transcriptome profiling identifies transcription factors and key homologs involved in seed dormancy and germination regulation of Chenopodium quinoa. Plant Physiology and Biochemistry 151, 443–456. doi:10.1016/j.plaphy.2020.03.050.

Xiong, R., Wang, Y., Wu, H., Ma, Y., Jiang, W. and Ma, X. (2018) Seed treatments alleviate dormancy of field bindweed (Convolvulus arvensis L.). Weed Technology 32, 564–569. doi:10.1017/wet.2018.46.

Zhou, Z.-Q., Bao, W.-K. and Wu, N. (2009) Dormancy and germination in Rosa multibracteata Hemsl. & E. H. Wilson. Scientia Horticulturae 119, 434–441. doi:https://doi.org/10.1016/j.scienta.2008.08.017.

